# Estrogen-Related Receptor is Required in Adult *Drosophila* Females for Germline Stem Cell Maintenance

**DOI:** 10.1101/2025.01.29.635514

**Authors:** Anna B. Zike, Madison G. Abel, Sophie A. Fleck, Emily D. DeWitt, Lesley N. Weaver

## Abstract

Stem cell self-renewal and proper tissue function rely on conserved metabolic regulators to balance energy production with inter-organ metabolic trafficking. The estrogen-related receptor (ERR) subfamily of orphan nuclear receptors are major transcriptional regulators of metabolism. In mammals, ERRs have roles in regulating mitochondrial biosynthesis, lipid metabolism, as well as stem cell maintenance. The sole *Drosophila ERR* ortholog promotes larval growth by establishing a metabolic state during the latter half of embryogenesis. In addition, *ERR* is required in adult *Drosophila* males to coordinate glycolytic metabolism with lipid synthesis and within the testis to regulate spermatogenesis gene expression and fertility. Despite extensive work characterizing of the role of *ERR* in *Drosophila* metabolism, whether *ERR* has a conserved requirement in regulating stem cell behavior has been understudied. To determine whether ERR regulates stem cell activity in *Drosophila*, we used the established adult female germline stem cell (GSC) lineage as a model. We found that whole-body *ERR* knockout in adult females using conditional heat shock-driven FLP-FRT recombination significantly reduces egg production and decreases GSC number. In addition, we found that ERR activity is required cell-autonomously in the adult female germline for maintenance of GSCs; whereas ERR regulation of GSCs is independent of its activity in adult female adipocytes. Our results highlight an ancient and conserved role for *ERRs* in the regulation of stem cell self-renewal.

**SUMMARY:** Tissue function and regeneration are dependent on stem cell populations, which relies on the coordination of energy homeostasis between multiple tissues within an organism. Zike *et al* report that the estrogen-related receptor (ERR) is required in adult *Drosophila* females to maintain germline stem cell number in the ovary to promote egg production. Furthermore, ERR regulation of stem cell maintenance is dependent on its cell autonomous role in the germline, but not through adipocyte-specific ERR activity. These findings demonstrate that the sole *Drosophila ERR* has a conserved role in regulating stem cell lineages (similar as the mammalian *ERRβ* ortholog) and can be used as a model to understand regulation of stem cell lineages as well as metabolism.

## INTRODUCTION

Adult tissue homeostasis is dependent on the balance of stem cell self-renewal and differentiation, which is largely governed by energy metabolism within an organism (Ghosh-Choudhary *et al*. 2020; Calibasi-Kocal *et al*. 2021; Jackson and Finley 2024). Stem cells are sensitive to physiological changes, including nutrient deprivation and environmental conditions (Ables *et al*. 2012). For example, a western diet of high fat and sugar increases mammalian intestinal stem cell proliferation and differentiation (Aliluev *et al*. 2021); whereas supplementation of nicotinamide adenine dinucleotide (NAD+) improves muscle stem cell function in muscular dystrophy mouse models (Zhang *et al*. 2016). Disruption of stem cell homeostasis can result in tissue dysfunction and diseases, including cancer (Reya *et al*. 2001; Shackleton 2010). However, the mechanisms by which the energetic need within an organism is balanced between multiple tissues to support stem cell lineages are not completely understood.

Nuclear receptors are transcription factors that act as physiological sensors to fine-tune the transcriptional landscape of a cell to maintain homeostasis across metabolically intensive tissues (Francis *et al*. 2003; King-Jones and Thummel 2005). Members of the estrogen-related receptor (ERR) subfamily of nuclear receptors are central regulators of energy homeostasis (Huss *et al*. 2015; Misra *et al*. 2017). In mammals, ERRα and ERRγ are required for mitochondrial biogenesis and lipid metabolism (Luo *et al*. 2003; Schreiber *et al*. 2004; Audet-Walsh and Giguere 2015; Fan *et al*. 2018; Fox *et al*. 2022), and ERRα acts in concert with its co-activator PGC-1alpha to regulate the transcription of genes involved in oxidative phosphorylation (Kamei *et al*. 2003; Schreiber *et al*. 2003). In addition to their metabolic activity, mammalian ERRs have been shown to regulate stem cell identity through transcriptional activation of self-renewal genes. For example, ERRβ is required for the maintenance of pluripotent embryonic stem cells through transcriptional regulation of Oct4, Nanog, and Sox2 (Festuccia *et al*. 2012; Festuccia *et al*. 2017). Knockdown of ERRβ in porcine induced pluripotent stem cells decreases expression of self-renewal genes, resulting in premature differentiation (Yang *et al*. 2018). These studies suggest ERRs have evolved specialized roles to regulate energy homeostasis and maintain stem cell identity by transcriptional regulation of targets in a tissue- or cell type-specific manner.

The *Drosophila* genome contains one ortholog of the ERR subfamily, eliminating the gene redundancy that is present in the mammalian genome (King-Jones and Thummel 2005). During larval stages, *ERR* is required in adipocytes for carbohydrate metabolism and *ERR* null mutants are hyperglycemic, dying at the second instar larval stage (Tennessen *et al*. 2011). In addition, *ERR* is required in adipocytes during the non-feeding pupal stage to up-regulate glucose oxidation and lipogenesis, promoting a metabolic state that persists during adulthood in *Drosophila* males (Beebe *et al*. 2020). ERRs have also been implicated in regulating reproduction in insects. In *Drosophila* males, loss of *ERR* in the testis reduces fertility due to mis-regulation of genes involved in spermatogenesis (Misra *et al*. 2017). Similarly*, Aedes aegypti ERR* transcription is activated after a blood meal in an ecdysone-dependent manner to regulate carbohydrate metabolism and oogenesis (Geng *et al*. 2024). Loss of *ERR* in mosquitos decreases lipid accumulation, disrupts carbohydrate metabolism, and significantly decreases egg production (Geng *et al*. 2024). Although extensive work has shown conservation of *ERR*-dependent metabolic regulation from insects to mammals, whether an ancestral *ERR* ortholog has a dual function to also regulate stem cell lineages is understudied.

The adult *Drosophila* ovary is a widely tractable model to understand how changes in whole-body energy homeostasis regulate stem cell lineages (Drummond-Barbosa 2019). Each female has two ovaries that contain 15-20 chains of progressively developing egg chambers (called ovarioles, **Figure 1a**) that originate from a germline stem cell (GSC) located at the anterior end of the ovariole in the germarium (**Figure 1b**). Each germarium contains two-to-three GSCs that divide asymmetrically to self-renew and give rise to differentiating progeny (Drummond-Barbosa 2019). GSCs are maintained in the niche (which is primarily composed of somatic cap cells) via multiple mechanisms. E-cadherin mediates GSC adhesion to the niche (Song and Xie 2002), which ensures physical proximity for GSCs to receive Bone Morphogenic Protein (BMP) signals that prevent differentiation (Xie and Spradling 1998). In addition to signaling from the niche, GSC self-renewal is dependent on intrinsic nuclear receptor activity. For example, the heterodimeric complex composed of EcR and Usp is required directly in GSCs to regulate their maintenance and proliferation (Ables and Drummond-Barbosa 2010). Furthermore, nuclear receptors act in somatic tissues outside of the ovary to indirectly influence GSC number. The nuclear receptor Seven up is required in adipocytes to regulate GSC maintenance (Weaver and Drummond-Barbosa 2019), whereas Hr4 activity in muscle indirectly maintains GSCs in the niche (Weaver and Drummond-Barbosa 2021). These studies suggest multiple nuclear receptors can coordinate their activities within the germline and in peripheral somatic tissues to control GSC self-renewal.

**Figure 1.**
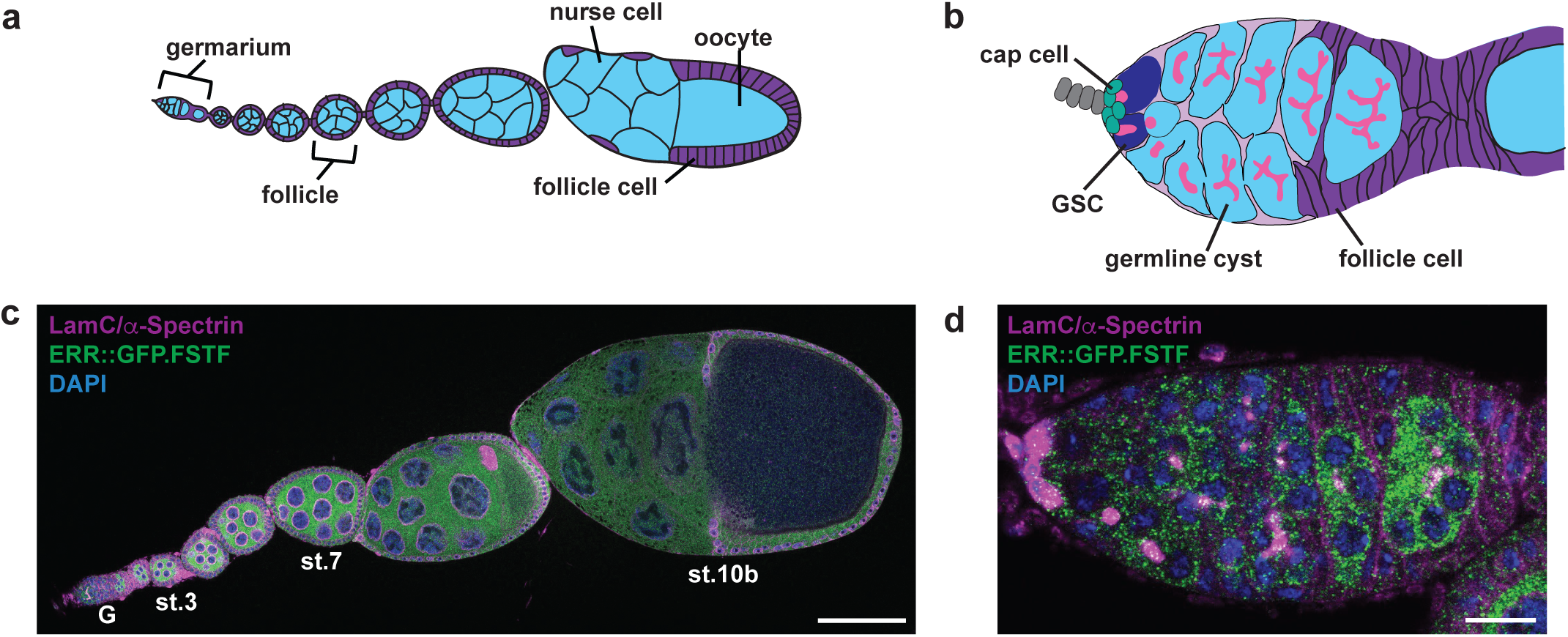
ERR is expressed in the germline of adult females. **(a)** The adult *Drosophila* ovary is composed of 15-20 ovarioles that contain progressively developing egg chambers. Each egg chamber consists of a germline cyst (15 nurse cells and one oocyte; blue) that is surrounded by somatic follicle cells (purple), which is produced in the anterior germarium. **(b)** Each germarium contains 2-to-3 germline stem cells (GSCs, dark blue) that are maintained by a somatic niche that is primarily composed of cap cells (green). GSCs divide to self-renew and give rise to a cystoblast that will undergo four mitoses with incomplete cytokinesis, generating a germline cyst. GSC and germline cysts are identified by the morphology of the fusome (pink), an ER-like organelle that changes morphology with differentiation. Germline cysts are surrounded by somatic follicle cells (purple) to form a follicle that ultimately buds from the germarium. **(c,d)** Ovariole **(c)** and germaria **(d)** from *ERR::GFP.FSTF* females showing the expression pattern of ERR::GFP.FSTF based on anti-GFP staining (green). α-Spectrin (magenta), fusome; LamC (magenta), nuclear lamina; DAPI (blue), DNA. Scale bar for (c), 100 µm; scale bar for (d), 10 µm. The germarium (G) and egg chamber stages are labeled in panel (c).

In this study we investigated the role for whole-body ERR activity in regulating the adult female *Drosophila* GSC lineage. Using conditional knockout alleles, we found that *ERR* is required in adult females for activation of known glycolytic and pentose phosphate pathway targets, egg production, and maintaining GSC number. We also found that ERR activity is required cell-autonomously in the germline to control GSC maintenance. Furthermore, we determined that ERR-dependent metabolic activity in adult female adipocytes is not required for oogenesis, although it is likely that ERR function in other somatic tissues is required for GSC maintenance. Our results highlight an ancient and conserved role for ERR in the regulation of stem cell self-renewal in adult *Drosophila* females.

## MATERIALS AND METHODS

### Drosophila strains and culture conditions

*Drosophila* stocks were maintained on BDSC cornmeal food (15.9 g/L inactive yeast, 9.2 g/L soy flour, 67.1 g/L yellow cornmeal, 5.3 g/L agar, 70.6 g/L light corn syrup, 0.059 M propionic acid) at 22-25°C. The previously described *ERR* alleles were used, including *dERR^Cond.19-4^*[RRID:BDSC_91656, (Beebe *et al*. 2020)], *dERR^cond.2^* (Beebe *et al*. 2020), *ERR^1^* [RRID:BDSC_83688, (Tennessen *et al*. 2011)], and *ERR^2^*[RRID:BDSC_83689, (Tennessen *et al*. 2011)]. In the *ERR^cond.19-4^* line, the endogenous *ERR* locus is flanked by two FRTs in combination with the *ERR^2^* mutant allele, removing the majority of the *ERR* coding region in response to heat shock to generate an *ERR* null animal. In the *ERR^cond.2^*construct, two FRTs are positioned around a fluorescently tagged *ERR* BAC transgene (*ERR::GFP.FSTF*) that is trans to *ERR^1^* and *ERR^2^*mutant alleles, resulting in an *ERR^1/2^* mutant animal upon heat shock. These constructs allow for animals to develop normally prior to adulthood when *ERR* removal is induced.

The previously described *tub-Gal80^ts^* [RRID:BDSC_7108, (McGuire *et al*. 2003)] and *3.1Lsp2-Gal4* [RRID:BDSC_98128, (Lazareva *et al*. 2007)] were used for adult, adipocyte-specific manipulations. The *w^1118^*allele (RRID:BDSC_5905) and *w^1118^; PBac{ERR-GFP.FSTF}VK00037* (RRID:BDSC_38638) line were obtained from the Bloomington *Drosophila* Stock Center (BDSC, bdsc.indiana.edu). The *UAS-mCas9* line was obtained from the Vienna *Drosophila* Resource Center (VDRC, https://shop.vbc.ac.at/vdrc_store/). Lines carrying multiple genetic elements were generated by standard crosses. Balancer chromosomes, *FRT* and *FLP* strains, and other genetic elements are described on Flybase (www.flybase.com).

### Quantification of egg laying and embryo hatching

The number of eggs laid were measured as previously described (Weaver and Drummond-Barbosa 2019) by maintaining five *ERR* conditional mutant or control females mated with five *w^1118^*males in perforated plastic bottles capped with molasses/agar plates smeared with active yeast paste. Molasses/agar plates were changed twice daily. The number of laid eggs per day was counted in five replicates per genotype and the results were subjected to a paired Student’s *t*-test.

Egg hatching was measured by transferring at least 20 eggs from molasses/agar plates to fresh molasses/agar plates containing active yeast paste in the center. The number of eggs that hatched were counted 24 hours after the transfer in five replicates per genotype. The results were subjected to a paired Student’s *t*-test.

### Generation of UAS-ERR 4x-sgRNA transgenes

sgRNA sequences targeting *ERR* were identified using the flyCRISPR Target Finder (http://targetfinder.flycrispr.neuro.brown.edu/) and selected sequences have no predicted off-target sites in the *Drosophila melanogaster* genome. A total of eight sgRNAs were used for the two transgene constructs (**Figure S6a; Table S2**), targeting unique regions within the *ERR* locus. Cloning of sgRNAs into the pCDF5 vector (Port and Bullock 2016) was carried out by GenScript and each 4x-sgRNA was verified by sequencing. Transgenesis was performed with the *PhiC31/attP/attB* system and plasmids were inserted at landing site (*P{ y[+]-attP-3B}VK00033*) on the third chromosome. Microinjection of plasmids was performed by Rainbow Transgenic Fly, Inc (Camarillo, CA).

### Adipocyte-specific ERR clustered regularly interspaced short palindromic repeats (CRISPR) mutagenesis

Females containing the *UAS-mCas9/cyo; UAS-4xgRNA/MKRS* of interest against *ERR* in combination with *y w; tubGal80^ts^/cyo; 3.1Lsp2/TM3* were raised at 18°C to block Gal4 activity (and therefore, CRISPR mutagenesis) during development. Zero-to-2-day-old females (with *Oregon-R* males) were maintained at 18°C for three additional days and subsequently switched to 29°C for CRISPR induction. *UAS-mCas9* was used as a negative control. For all conditions, medium was supplemented with wet active yeast paste.

### Clonal mosaic analysis

Females of genotype *hs-FLP/+; His2Av-mRFP FRT2A/ERR* FRT2A* or *hs-FLP/+; PBac{ERR-GFP.FSTF}/+; His2Av-mRFP FRT2A/ERR* FRT2A* (where *ERR\** denotes *ERR* null alleles or *wild-type*) were generated by standard crosses. Zero-to-two day old females were heat shocked at 37°C twice daily to induce *flippase (FLP)/FLP recognition target (FRT)*-mediated mitotic recombination, as previously described (Laws and Drummond-Barbosa 2015). After the final heat shock, females were maintained at 25°C for 4, 8, or 12 days on medium supplemented with dry yeast. Females were transferred to media supplemented with wet yeast paste two days prior to dissection. *ERR\** homozygous clones were recognized based on their loss of mRFP expression. We determined GSC loss by calculating the percentage of germaria containing mRFP-negative cystoblasts and/or cysts that had lost the mRFP-negative progenitor GSC (Laws and Drummond-Barbosa 2015). Three independent experiments were performed, and statistical significance was calculated using a paired Student’s *t*-test.

### Adult female tissue immunostaining and fluorescence microscopy

Adult female ovaries were dissected in unsupplemented Grace’s Insect Medium (Gibco), fixed, and washed as described (Weaver and Drummond-Barbosa 2019). Samples were blocked at least 3 hours at room temperature in 5% normal goat serum (NGS; Jackson ImmunoResearch) plus 5% bovine serum albumim (BSA; Sigma) in phosphate-buffered saline (PBS; 10 mM NaH_2_PO_4_/NaHPO_4_, 175 mM NaCl, pH 7.4) containing 0.1% Triton-X100 (PBST), and incubated overnight at 4°C in primary antibodies diluted in blocking solution as follows: mouse monoclonal anti-α-Spectrin concentrate (3A9-c) (DSHB; Developmental Studies Hybridoma Bank, 3 µg/ml); mouse monoclonal anti-Lamin C (LC28.26) (DSHB, 0.8 µg/ml); rat monoclonal anti-E-Cadherin concentrate (DCAD2-c) (DSHB, 0.37 µg/ml); and rabbit polyclonal anti-Vasa (Boster Bio, 1 µg/ml). Samples were washed in PBST and incubated for 2 hours at room temperature in 1:200 Alexa Fluor 488- or 568-conjugated goat species-specific secondary antibodies (Molecular Probes) in blocking solution. Stained samples were washed, mounted in Vectashield containing 1.5 µg/ml 4’,6-diamidino-2-phenylindole (DAPI) (Vector Laboratories), and imaged using a Nikon SP8 confocal microscope.

GSCs and cap cells were identified as described (Weaver and Drummond-Barbosa 2019). For analysis of *ERR* conditional knockout assays, a Student’s *t*-test was used to calculate the statistical significance of any differences among the control (no heat shock) and experimental conditions (heat shock) from at least three independent experiments. For adipocyte-specific CRISPR mutagenesis experiments, two-way ANOVA with interaction (GraphPad Prism) was used to calculate the statistical significance of differences among genotypes in the rate of GSC loss from at least three independent experiments, as described (Armstrong *et al*. 2014).

For Enolase (Eno) protein level analysis, ovaries were dissected in Grace’s Insect Medium and fixed for 20 minutes in 4% formaldehyde, 1X PBS, and 0.1% Tween 20. Ovaries were washed once for 10 minutes in solution containing 1X PBS and 0.1% Tween. Ovaries were blocked at room temperature for one hour in blocking solution containing 1X PBS, 10% BSA, and 0.1% Tween 20 and incubated over night at 4°C in Enolase primary antibody (1:100, Santa Cruz Biotechnology) diluted in 1X PBS, 0.1% Tween 20, and 1% BSA. Samples were washed three times and subsequently incubated in secondary antibody for four hours at room temperature. Samples were processed for the remainder of immunofluorescence as described above. The densitometric mean of the cytoplasm from individual GSCs was measured using FIJI (https://fiji.sc/) as previously described (Rojas-Rios *et al*. 2024).

For pMad immunofluorescence, ovaries were dissected in Grace’s Insect Medium and fixed for 15 minutes in 4% formaldehyde. Ovaries were washed in solution containing 1X PBS, 0.1% Triton-X100, and 0.05% Tween 20. Ovaries were blocked for at least 3 hours in 1X PBS, 0.1% Triton-X100, 0.05% Tween 20, and 1% BSA, and incubated over two nights at 4°C in primary antibody in blocking solution. Primary antibodies used include rabbit polyclonal anti-pS423/S425 Smad3 (1:100, Abcam), mouse monoclonal anti-α-Spectrin concentrate (3A9-c) (DSHB; Developmental Studies Hybridoma Bank, 3 µg/ml); and mouse monoclonal anti-Lamin C (LC28.26) (DSHB, 0.8 µg/ml). Samples were washed three times and processed for the remainder of immunofluorescence as described above. The densitometric mean of individual GSC nuclei was measured from optical sections containing the largest nuclear diameter (visualized by DAPI) using FIJI as previously described (Armstrong *et al*. 2014).

For E-Cadherin quantification, the total densiometric value from maximum intensity projections around cap cells (identified using LamC staining) was measured with FIJI (Weaver and Drummond-Barbosa 2018), which is a well-established method to determine whether adhesion to the GSC niche is compromised (Hsu and Drummond-Barbosa 2011). Densitometric data for Eno, pMad, and E-Cadherin analysis were subjected to the Mann-Whitney *U*-test from at least three independent experiments.

For adipocyte morphology and lipid droplet visualization, fixed and washed fat bodies attached to abdominal carcasses were incubated with 2.5X Alexa Fluor 488-conjugated phalloidin (Molecular Probes) in PBST for 20 min, rinsed, and washed three times for 15 min each in PBST. Samples were stored in Vectashield plus DAPI (Vector Laboratories) containing 25 ng/ml Nile Red dye, and fat bodies were scraped off the abdominal carcass before mounting and imaging using a Nikon SP8 confocal. The largest cell area of each adipocyte (based on phalloidin staining) used to measure adipocyte area; whereas the longest width of each nucleus per adipocyte (based on DAPI) determined nuclear diameter. Each measurement was performed using FIJI. Three independent experiments were performed, and statistical analysis was performed using a Student’s *t* test.

### Analysis of GSC and follicle cell proliferation, death of early cysts, and dying vitellogenic follicles

For EdU incorporation analysis, intact ovaries were incubated at room temperature for one hour in 100 µM EdU (Life Technologies) diluted in room temperature Grace’s media. After incubation, ovaries were washed, ovarioles were separated, and fixed as described above. After incubation with primary antibodies, samples were subjected to the Click-iT reaction according to the manufacturer’s instructions (Life Technologies) for 30 minutes at room temperature. GSC S-phase and M-phase cell cycle percentages were measured by calculating the fraction of EdU- or phospho-histone H3-positive GSCs as a percentage of the total number of GSCs analyzed per genotype. Follicle cell proliferation was calculated by counting the average percentage of phospho-histone H3-positive follicle cells and EdU-positive follicle cells from stage 4-6 egg chambers. Single confocal plane images of the top and bottom of mounted ovarioles were used for analysis as previously described (Laws and Drummond-Barbosa 2016).

### RNA isolation, reverse transcriptase (RT)-qualitative polymerase chain reaction (RT-qPCR)

Ten whole adult females of each control or *ERR* conditional knockout genotype were lysed in 500 µl lysis buffer from the RNAqueous-4PCR DNA-free RNA isolation for RT-PCR kit (Invitrogen). RNA was extracted from all samples following manufacturer’s instructions from three independent experiments.

Adult adipocyte RNA extraction was performed as previously described (Weaver and DDB 2019). Briefly, abdominal carcasses from 100 females of each genotype were dissected in Grace’s medium supplemented with 10% fetal bovine serum. Fat bodies were removed from the abdominal carcass with 500 µl dissociation buffer [0.5% Trypsin (Sigma) plus 1 mg/ml collagenase (Sigma) in 1X PBS] per 50 carcasses for 30 minutes at room temperature. 500 µl of Grace’s media plus 10% FBS was added to stop the enzymatic reaction, and the supernatant was placed in a new tube. Carcasses were rinsed with Grace’s media plus 10% FBS and the two supernatants per genotype were combined. Cells were centrifuged at 3.3 rpm for 5 minutes at room temperature. After removal of supernatant, cells were flash frozen in liquid nitrogen and stored at −80°C until RNA extraction. Cells were lysed in 250 µl lysis buffer from the RNAqueous-4PCR DNA-free RNA isolation for RT-PCR kit (Invitrogen), and RNA was extracted following the manufacturer’s instructions from three independent experiments.

cDNA was synthesized from 500 ng of total RNA extracted from whole females or female adipocytes described above using Superscript II Reverse Transcriptase (Thermo Fisher Scientific) according to the manufacturer’s instructions. **Table S1** lists all primers used in this study. *Rp49* and *Act5C* transcripts were used as controls. PowerUp SYBR Green Master Mix (Thermo Fisher Scientific) was used for qRT-PCR. Reactions for three independent biological replicates were performed in triplicate using LightCycler 96 (Roche). The fluorescence amplification threshold was determined by LightCycler 96 software, and ddCT were calculated using Microsoft Excel. Fold change of transcript levels was calculated in Excel as described (Taylor *et al*. 2019).

### Fly liquid-food interaction counter (FLIC) assay

The FLIC system was used as previously described (Ro *et al*. 2014) to determine whether *ERR* knockout disrupted feeding behavior. FLIC *Drosophila* Feeding Monitors (DFMs, Sable Systems International, models DFMV2 and DFMV3) were used in the single choice configuration. Each chamber was loaded with liquid food solution that contained 4% sucrose (m/v) and 1.5% yeast extract (m/v). 10-day-old females were aspirated into the DFM chambers and feeding behavior was measured for 24 hours. Each FLIC experiment contains pooled data from at least 30 females for each genotype. FLIC data were analyzed using a previously described custom R code (Ro *et al*. 2014). Default thresholds were used for analysis except the following: minimum feeding threshold = 10, tasting threshold = (0,10). Females that did not participate (i.e., returned zero values) or produced extreme outliers were excluded from analysis. Data were subjected to a Mann-Whitney *U*-test.

## RESULTS

### ERR is expressed in the adult Drosophila ovary

To determine whether ERR activity is required in the adult female GSC lineage, we first assayed the expression and localization pattern of ERR protein in the ovary using the fluorescently tagged ERR BAC transgene (*ERR::GFP.FSTF*) in which *ERR* is expressed from its endogenous locus. We found that ERR::GFP.FSTF is expressed throughout the ovary, with expression starting in the germarium and continuing through stage 10 (**Figure 1c,d**). Interestingly, ERR::GFP.FSTF is localized to the cytoplasm of germ cells instead of the nucleus. Although the ERR::GFP.FSTF transgene contains a multiple reporter tags that may impede proper ERR cellular targeting, the cytoplasmic localization observed is consistent with sequestration of other nuclear receptors in the cytosol by molecular chaperones until they activated (Levin and Hammes 2016). These results suggest that ERR is expressed in germline and may have non-nuclear functions to regulate the GSC lineage.

### Whole-body ERR activity is required for egg production in adult females

ERR regulates energy homeostasis in multiple *Drosophila* tissues (Tennessen *et al*. 2011; Beebe *et al*. 2020; Holcombe and Weavers 2023; Fasteen *et al*. 2024), and loss of *ERR* during larval stages results in lethality prior to adulthood (Tennessen *et al*. 2011). Therefore, we utilized previously described conditional knockout alleles of *ERR* [hereafter referred to as *ERR^cond.19-4^* and *ERR^cond.2^*(Beebe *et al*. 2020)] to assess the requirement of whole-body ERR activity during oogenesis in adults, which allow for animals to develop normally prior to deletion of the *ERR* locus in adults.

To determine the role of *ERR* in regulating oogenesis specifically during adulthood, we subjected zero-to-two day old *ERR^cond.19-4^*or *ERR^cond.2^* adult females to sequential heat shock treatments at 37°C separated by 24 hours and compared the number of eggs laid to their respective no heat shock sibling control for 20 days (**Figure 2a,b**). Conditional knockout of *ERR* using the *ERR^cond.19-4^* experimental females resulted in fewer eggs laid relative to control (**Figure 2a**), however the differences were not statistically significant. Conversely, *ERR^cond.2^*conditional knockout females laid significantly fewer eggs relative to their sibling controls (**Figure 2b**), suggesting the *ERR^cond.2^* conditional knockout condition may be stronger than *ERR^cond.19-4^*. For both conditional knockout lines, there were no significant differences in embryonic viability (**Figure S1a,b**), indicating that there is not a maternal requirement for *ERR* in resulting progeny. These results suggest that *ERR* is required for egg production in adult females.

**Figure 2.**
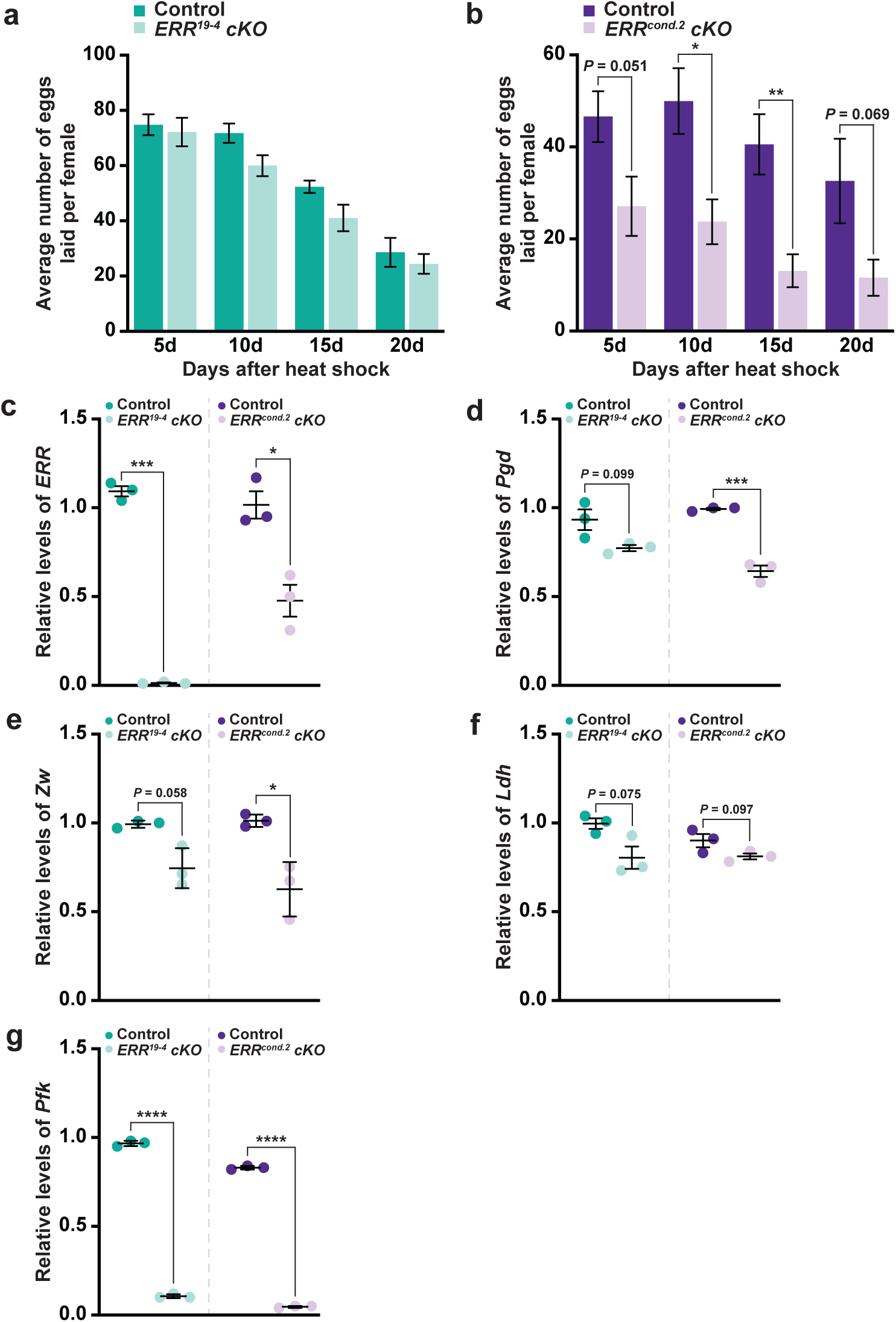
*ERR* is required in adult females for egg production and metabolic enzyme transcription. **(a,b)** The average number of eggs laid by adult females for *ERR^19-4^* **(a)** or *ERR^cond.2^* **(b)** and their respective controls at 5, 10, 15, and 20 days after the final heat shock. Data is shown as the average ± standard error of the mean (SEM). \**P* < 0.05, \*\**P* < 0.01 (paired, two-tailed Student’s *t*-test). Five biological replicates were performed for each genotype. **(c-g)** Quantitative reverse-transcriptase polymerase chain reaction (qRT-PCR) analysis of *ERR* **(c)**, *Pgd* **(d)**, *Zw* **(e)**, *Ldh* **(f)**, and *Pfk* **(g)** transcripts from whole-body females in which *ERR* was conditionally knocked out using the *ERR^19-4^* or *ERR^cond.2^* conditional knockout alleles. Females were collected 7 days after the final heat shock and compared to their respective control. Experiments were performed in triplicate. Data is shown as average ± standard error of the mean (SEM). \**P* < 0.05, \*\*\**P* < 0.001, \*\*\*\**P* < 0.0001 (paired, two-tailed Student’s *t*-test).

To determine the relative strength of the different conditional knockout lines used, we isolated total RNA from control and *ERR* conditional knockout adult females 7 days after heat shock and performed quantitative reverse-transcriptase polymerase chain reaction (qRT-PCR) to quantify the transcript levels of *ERR* and known ERR transcriptional target genes (**Figure 2c-g**). Relative to control, *ERR* transcripts using *ERR^cond.19-4^* were reduced to negligible levels. Surprisingly, *ERR^cond.2^* females retained ∼50% of *ERR* transcripts relative to control (**Figure 2c**). It is likely that *ERR* transcripts are still detected in the *ERR^cond.2^* line due to the presence of the *ERR^1^* allele, which contains point mutations in exons 2 and 3, yet results in a full-length *ERR* transcript (Tennessen *et al*. 2011). Consistent with this idea, transcript levels of pentose phosphate pathway targets *Pgd* and *Zw* were significantly decreased in the *ERR^cond.2^*knockout (with a moderate, albeit not significant, decrease in the *ERR^cond.19-4^*line) (**Figure 2d,e**). ERR glycolytic targets were also decreased in both conditional knockout lines (**Figure 2f,g**), with the most significant reductions from the *ERR^cond.2^* line. These results suggest that while both *ERR* conditional knockout females reduce the levels of *ERR* and known targets, *ERR^cond.2^*is a stronger genetic manipulation.

### ERR is required in adult females to maintain GSC number

Continuous production of mature oocytes is dependent on the maintenance and proliferation of the GSC population located in the germarium (Hsu *et al*. 2008). Because *ERRβ* is required in mammals for stem cell self-renewal (Festuccia *et al*. 2012), we hypothesized that loss of *ERR* in adult *Drosophila* females may influence the GSC lineage. We therefore assayed whether defects in egg production were due to a decrease in GSC number by comparing the number of GSCs in *ERR* conditional knockout females relative to control over 15 days (**Figure 3**). Relative to control, *ERR* conditional knockout germaria have no obvious defects in overall morphology (**Figure 3a,b**). However, when we quantified GSC number in control and experimental females by counting the cells with an anteriorly localized fusome attached to the cap cells, both *ERR^cond.19-4^* and *ERR^cond.2^* conditional knockouts resulted in significantly fewer GSCs at each timepoint relative to the sibling no heat shock controls (**Figure 3c,d**). These results suggest that *ERR* is required in at least one tissue of the adult female to maintain GSCs in the niche. We would note, however, that the decrease in GSC number is not due to a change in the niche size, which regulates GSC number (Hsu and Drummond-Barbosa 2009), as the number of cap cells is not significantly different between control and experimental animals (**Figure S2**). Overall, our findings suggest that the ERR requirement to regulate stem cell lineages is conserved from insects to mammals.

**Figure 3.**
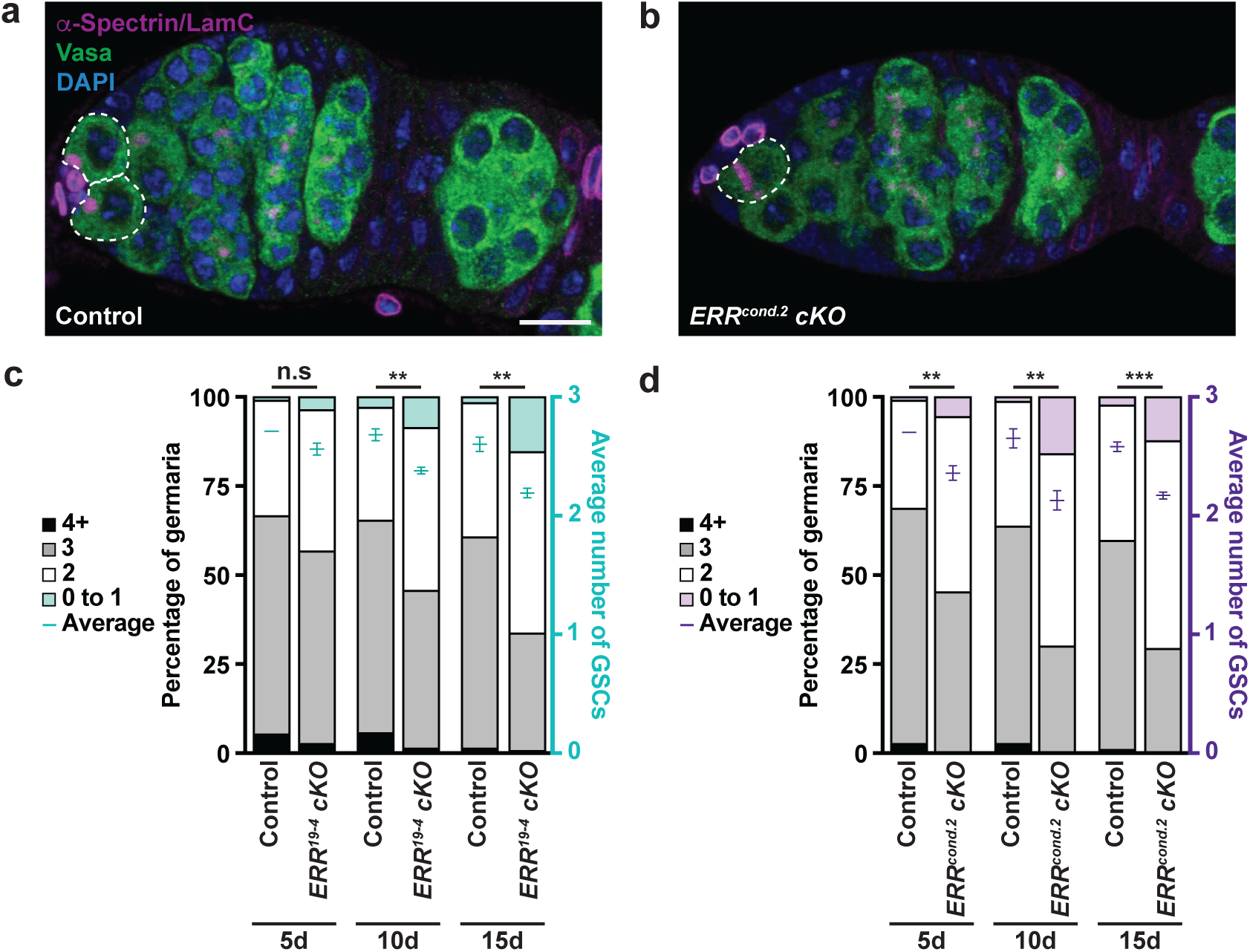
GSC number is decreased in *ERR* conditional knockout females. **(a,b)** Germaria from females at 15 days in control **(a)** or *ERR^cond.2^* **(b)** females 15 days after the final heat shock. Vasa, germ cells (green); α-Spectrin, fusome (magenta); LamC, nuclear lamina (magenta); DAPI, DNA (blue). GSCs are outlined. **(c,d)** Bars representing the percentage of germaria containing zero, one, two, three, or four-or-more GSCs at 5, 10, and 15 days after the final heat shock of females from *ERR^19-4^* **(c)** and *ERR^cond.2^* **(d)** conditional knockout females relative to their respective controls. The average GSC numbers are shown as mean ± standard error of the mean (SEM) is plotted on the right y-axis. 300 germaria were analyzed from three biological replicates. n.s., not significant, \*\**P* < 0.01, \*\*\**P* < 0.001 (paired, two-tailed Student’s *t*-test).

### ERR does not regulate E-Cadherin levels or BMP signaling

Because the decrease in GSC number is not due a reduction in the GSC niche, we next asked whether known regulators of GSC maintenance were affected in *ERR* conditional knockout females. GSC anchorage to cap cells is mediated by the adhesion protein, E-Cadherin, which is essential for GSC maintenance (Song and Xie 2002); whereas bone morphogenic protein (BMP) signaling is required to maintain stem cell identity and prevent differentiation (Xie and Spradling 1998). To determine whether the relative levels of E-Cadherin were altered, we measured the densiometric intensity of E-Cadherin protein in *ERR^19-4^* and *ERR^cond.2^* knockout females 10 days after heat shock and found there was not a significant decrease in E-Cadherin levels relative to their respective controls (**Figure S3a,b**). In addition, we measured the intensity of nuclear-localized phosphorylated Mad (pMad, a BMP signaling reporter, (Kai and Spradling 2003) in females 10 days after heat shock, and determined that the levels were statistically similar between control and *ERR* conditional knockout females (**Figure S3c,d**). These results suggest that the differences in GSC number when *ERR* is knocked out in the adult female is not a result of premature differentiation due to decreased adherence to the niche or a decrease in BMP signaling.

### Whole-body ERR activity is required for germline glycolytic enzyme expression

Stem cells are rewired to preferentially utilize glycolysis instead of oxidative phosphorylation to maintain stem cell identity (Folmes and Terzic 2016). It was recently shown that translational activation of glycolytic mRNAs in the adult female germline (including the ERR targets *Eno*, *pyk*, and *Ald*) are dependent on PIWI-interacting RNAs (piRNAs) and PIWI proteins (Rojas-Rios *et al*. 2024), and inhibition of glycolytic enzyme translation results in GSC loss (Rojas-Rios *et al*. 2024). To determine whether whole-body knockout of *ERR* decreases transcription, and therefore translation, of enzymes required for metabolic reprograming in GSCs, we measured protein levels of Enolase (Eno) in control and *ERR* conditional knockout females 10 days after heat shock (**Figure 4a,b**). Relative to control, *ERR* conditional knockout GSCs had a significant decrease in Eno levels, with *ERR* mutant GSC expression decreased by about 25% (**Figure 4c**). These results suggest that the decrease in GSCs in *ERR* conditional knockout females may be due to decreased glycolytic enzyme expression and failure to rewire GSCs for their maintenance.

**Figure 4.**
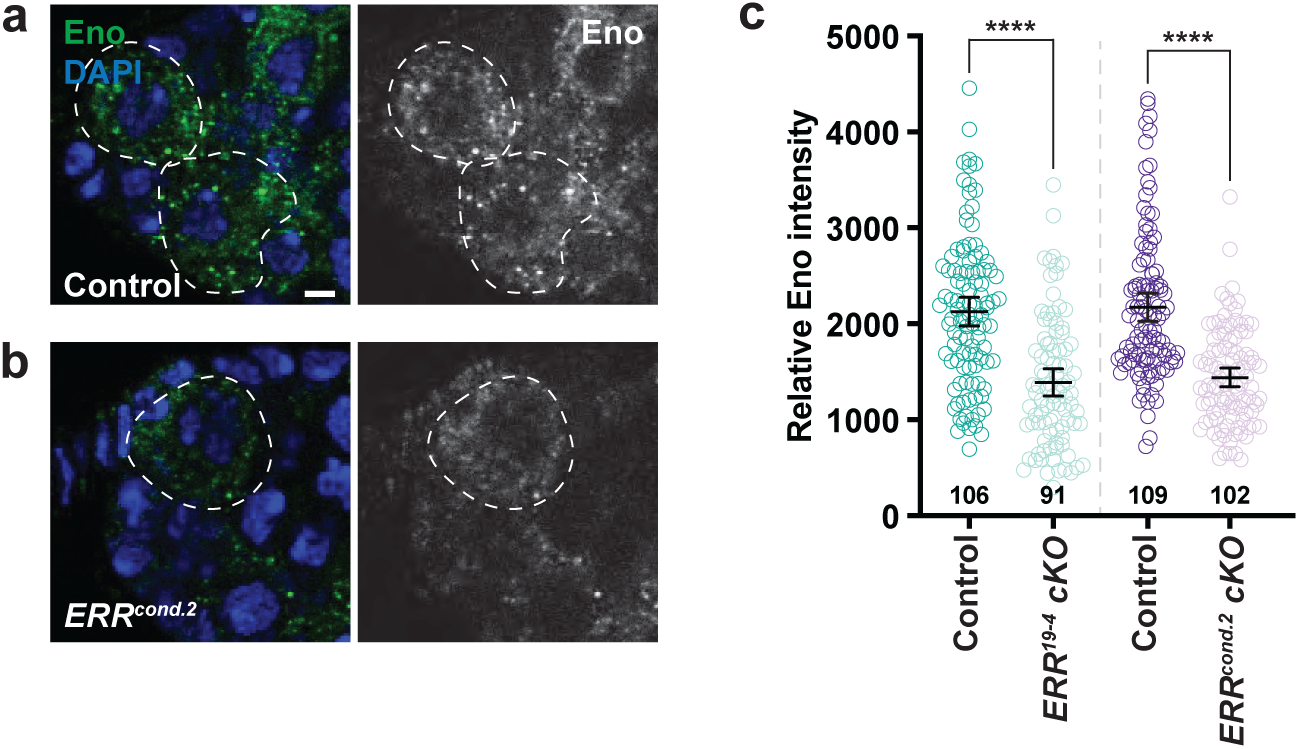
*ERR* conditional knockout in adult females reduces glycolytic enzyme expression in GSCs. **(a,b)** Anterior portion of germaria from control **(a)** and *ERR^cond.2^* mutant **(b)** females 10 days after heat shock showing Eno levels in GSCs. Eno (green); DAPI (blue), DNA. Scale bar, 2.5 µm. **(c)** Dot plot of Eno intensity per GSC for each genotype. The number of GSCs analyzed from three independent experiments is indicated in the graph. Data shown as mean ± 95% confidence interval for each genotype. \*\*\*\**P* < 0.0001; Mann-Whitney *U*-test.

### Loss of ERR in adult females does not influence eating behavior or adipocyte morphology

Defects in oogenesis could result from a decrease in food consumption (Gandara and Drummond-Barbosa 2022; Nunes and Drummond-Barbosa 2023) or severe perturbation of adipose tissue morphology (Matsuoka *et al*. 2017). To determine whether feeding behavior was altered in females lacking *ERR*, we used the Fly Liquid-Food Interaction Counter (FLIC; (Ro *et al*. 2014)) to assess their food consumption relative to controls over a 24-hour feeding period (**Figure S4a,b**). Compared to controls, *ERR^cond.19-4^* and *ERR^cond.2^* females had similar numbers of encounters with the liquid food and there was no change in the duration *ERR* conditional knockout females ate. Furthermore, when we assessed overall adipocyte morphology, we did not observe any difference in the adipocyte size or nuclear diameter in *ERR* conditional knockout females relative to controls (**Figure S4c-e**). These results suggest that changes in feeding behavior or gross alteration of adipocyte morphology do not account for the decrease in GSC maintenance.

### ERR is not required in adult females for GSC proliferation or early cyst survival

In addition to maintenance of GSCs, defects in fertility can also occur due to decreased GSC proliferation or death of early dividing and differentiating progeny (Drummond-Barbosa and Spradling 2001; Hsu *et al*. 2008). To determine whether GSC proliferation is altered when *ERR* is conditionally knocked out in adult females, we measured the percentage of GSCs that incorporated the thymidine analog, 5-ethynyl-2’-deoxyuridine (EdU, representing S-phase GSCs), and those that were labeled by phospho-histone H3 (pHH3, representing GSCs in mitosis). There was not a significant reduction in the GSC proliferation of *ERR^cond.19-4^* nor *ERR^cond.2^* conditional knockout females compared to their relative controls (**Figure S5a,b**), indicating that the decrease in egg production in *ERR* knockout females is not due to decreased GSC proliferation.

To determine whether whole-body ERR activity is required for survival of early differentiating progeny, we used a TUNEL-based assay that detects significant DNA fragmentation [ApopTag; (Drummond-Barbosa and Spradling 2001)] in *ERR* conditional knockout females 7 days after heat shock and compared the percentage of germaria with ApopTag positive germline cysts relative to control. Adult specific loss of *ERR* did not significantly increase the percentage of germaria containing dying germline cysts relative to non-heat shock controls (**Figure S5c**). These results suggest that neither GSC proliferation nor survival of differentiating progeny require ERR activity in adult females.

### Egg chamber growth and vitellogenic egg chamber survival are not dependent on whole-body ERR

In addition to the early stages of oogenesis that occur in the germarium, oogenesis is also dependent on the proper growth of egg chambers and oocyte yolk accumulation during vitellogenesis (Laws and Drummond-Barbosa 2017; Drummond-Barbosa 2019). Therefore, we tested whether whole-body ERR activity is required for these developmental processes. To assess egg chamber growth, we quantified the percentage of follicle cells in S-phase (based on EdU incorporation) or M-phase (based on pHH3 staining) from egg chambers in stages two through six in *ERR* conditional knockout females relative to control 7 days after heat shock (**Figure S5d,e**). The percentages of S-phase and M-phase proliferating follicle cells were not significantly different in *ERR* conditional knockout females compared to control. To determine whether *ERR* is required in the adult female for vitellogenesis, we counted the percentage of dying vitellogenic egg chambers based on pyknotic nuclei (visualized by DAPI). Both *ERR* conditional knockout and control females had negligible vitellogenic egg chamber death (zero-to-one percent for all experimental conditions). Collectively, these results indicate that *ERR* is not required in adult females for egg chamber growth or survival of vitellogenic follicles.

### Adipocyte ERR activity is not required in adult females for GSC maintenance

Our results thus far suggest that ERR activity is required in at least one adult female tissue or cell type to exclusively regulate GSC maintenance and no additional ovarian process. *ERR* is required in larval and adult male adipocytes for regulation of triglycerides (Tennessen *et al*. 2011; Beebe *et al*. 2020; Fasteen *et al*. 2024). Furthermore, nutrient-sensing and metabolic activity in the adipose tissue is required for multiple processes of oogenesis (Armstrong *et al*. 2014; Matsuoka *et al*. 2017; Armstrong and Drummond-Barbosa 2018; Weaver and Drummond-Barbosa 2019), making this an ideal candidate tissue to assess. We used the *ERR::GFP.FSTF* BAC transgene to assess ERR expression and localization in adult female adipocytes. Similarly as reported for larval adipocytes (Fasteen *et al*. 2024), ERR is localized to the nucleus of adult female adipocytes (**Figure 5a**). To determine whether adipocyte-specific ERR activity is required for GSC maintenance, we generated two *UAS-sgRNA* constructs that each contain 4 sgRNAs, collectively targeting eight distinct regions of the *ERR* locus (**Figure S6a; Table S2**). We then induced adult-specific CRISPR mutagenesis specifically in adipocytes using the previously described *3.1Lsp2-Gal4* (Lazareva *et al*. 2007) combined with *tub-Gal80^ts^*(Armstrong *et al*. 2014). Compared to *UAS-mCas9* control, CRISPR mutagenesis under control of each of the *UAS-sgRNA*s significantly decreased *ERR* transcripts (**Figure 5b**) and did not have any obvious morphological effects on the adipose tissue (**Figure S6b,c**). We found that adipocyte-specific *ERR* disruption did not significantly alter the rate of GSC loss relative to control (**Figure 5c, Figure S6d**). These results suggest that *ERR* is not required in adult female adipocytes to regulate GSC number. However, it is possible that ERR activity may be required in another metabolically-intensive tissue to regulate GSC self-renewal.

**Figure 5.**
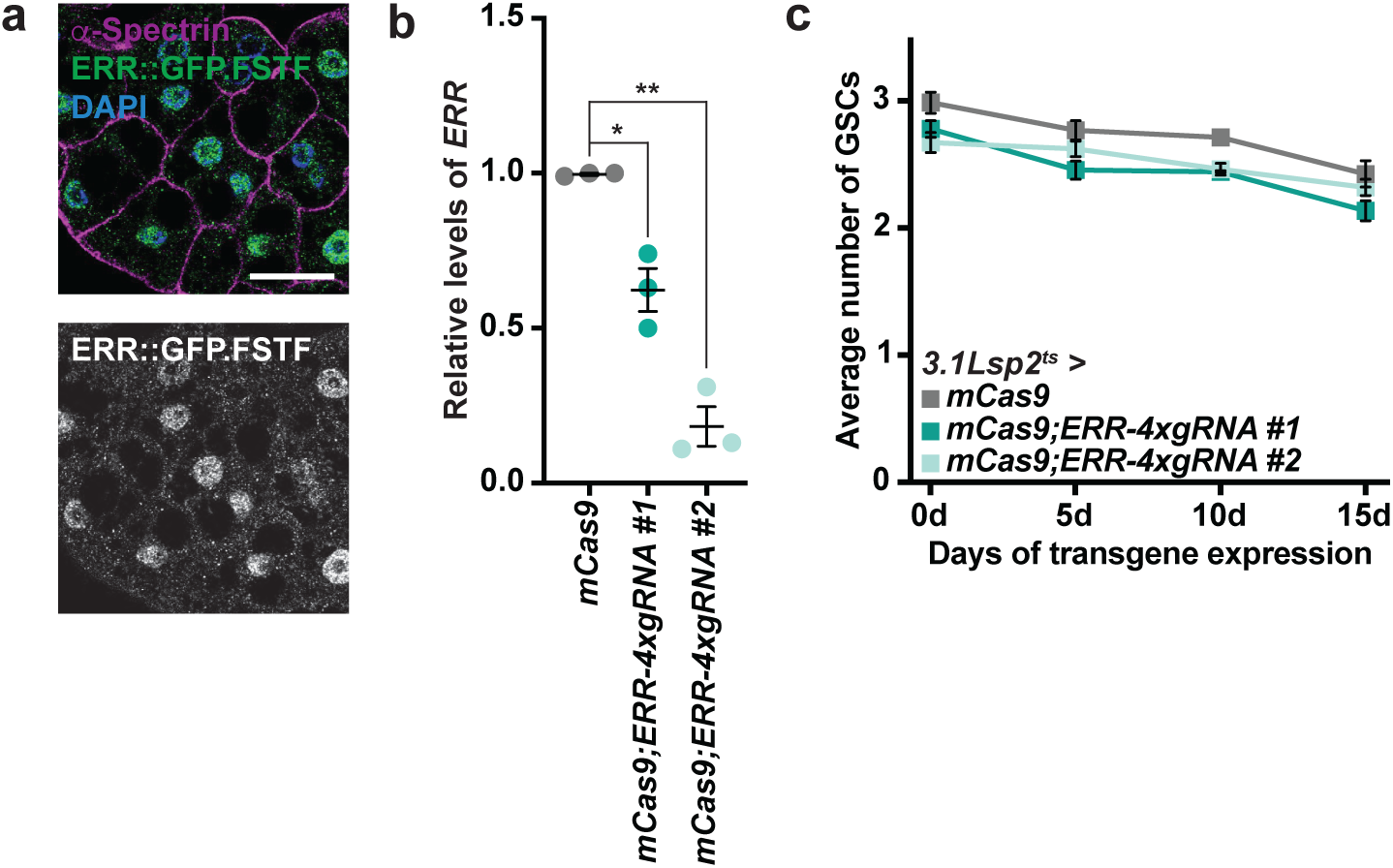
*ERR* is not required in adult female adipocytes for GSC maintenance. **(a)** Localization of ERR::GFP.FSTF (green) in adult female adipocytes. α-Spectrin (magenta), cell outline; DAPI (blue), DNA. Scale bar, 25 µm. **(b)** Quantitative reverse transcriptase polymerase chain reaction (qRT-PCR) analysis of *ERR* transcripts when *ERR* is knocked out specifically in adult female adipocytes using the *3.1Lsp2-Gal4* driver combined with *tubGal80^ts^*(*3.1Lsp2-Gal4^ts^*). Three independent experiments were performed. Data shown as mean ± SEM. \**P* < 0.05, \*\**P* < 0.01 (paired, two-tailed Student’s *t*-test). **(c)** The average number of GSCs over time in females with adult adipocyte-specific CRISPR using *UAS-mCas9* control or two *UAS-ERR* sgRNA constructs driven by *3.1Lsp2-Gal4^ts^*. Data shown as mean ± SEM. No statistically significant differences, two-way ANOVA with interaction.

### ERR is required in the germline for GSC maintenance

Because the *ERR* conditional knockout removes *ERR* from both the germline and somatic tissues, we also examined the role of ERR activity for GSC maintenance within the germline. To assess the autonomous role of *ERR* in the germline to regulate oogenesis, we used the previously described *ERR^1^* and *ERR^2^* mutant alleles (Tennessen *et al*. 2011) for genetic mosaic analysis and assessed GSC maintenance (**Figure 6**). In control mosaic females, in which both mRFP-positive and mRFP-negative GSCs are wild-type (**Figure 6a**), we observed a slight decrease in the number of mRFP-negative GSCs retained at the niche over time (**Figure 6c**). Conversely, *ERR^1^*and *ERR^2^* mutant GSCs were rapidly lost from the niche, which was indicated by the presence of mRFP-negative differentiating progeny without a respective mRFP-negative GSC (**Figure 6b,c**). By 8 days after heat shock, more than 23% of *ERR* mutant mosaic germaria of either genotype had lost GSCs compared to 7.84% in control (**Figure 6c,d**). We also found that the loss of GSCs were dependent on ERR activity, as the GSC loss events were partially rescued in the presence of the ERR::GFP.FSTF activity (**Figure 6c,d**). Taken together, our results suggest that *ERR* is required cell-autonomously to regulate GSC maintenance.

**Figure 6.**
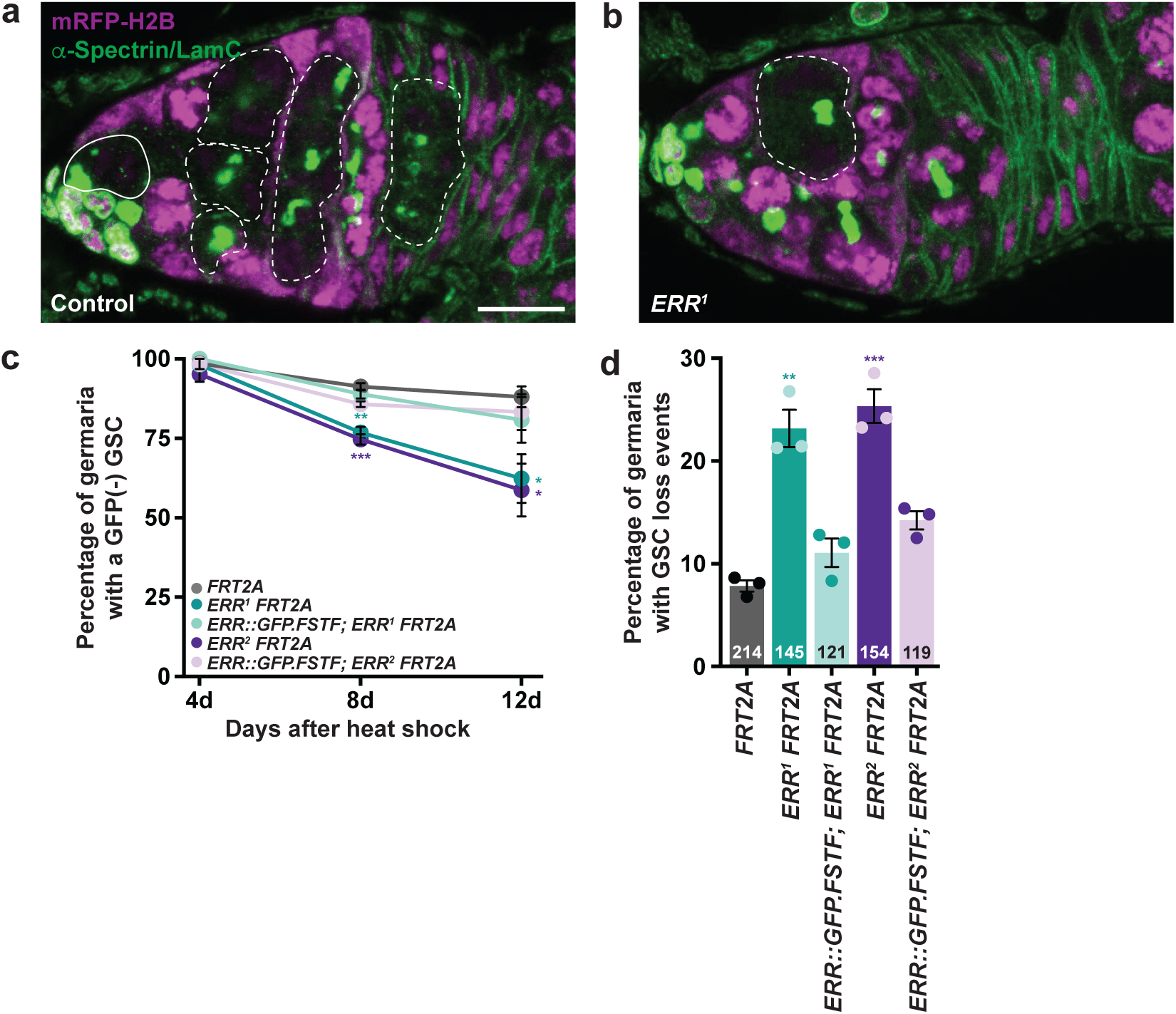
*ERR* is required in the germline for GSC maintenance. **(a,b)** Genetic mosaic germaria showing mRFP-negative control (from “mock” mosaics; **a**) or *ERR^1^*homozygous (**b**) cystoblasts and cysts (dashed lines) at 12 days after heat shock. The GSC negative mother is present in (**a**) (solid outline) from which mRFP-negative cysts derive in control mosaics, it is absent in (**b**) representing a GSC loss event. Scale bar for (**a**) and (**b**), 10 µm. **(c)** The percentage of total germaria with a GFP-negative GSC at 4, 8, and 12 days after heat shock used to measure GSC loss. At least 65 mosaic germaria were analyzed for each condition. Data is shown as the mean ± standard error of the mean (SEM) from three independent experiments. \**P* < 0.05, \*\**P* < 0.01, \*\*\**P* < 0.001 (paired, two-tailed Student’s *t*-test). **(d)** The percentage of mosaic germaria with a GSC loss event at 8 days after heat shock. This is the same 8 day after heat shock data represented in (**c**). The number of mosaic germaria analyzed from three biological replicates are indicated in each bar. Data is shown as mean ± SEM. \*\**P* < 0.01, \*\*\**P* < 0.001 (paired, two-tailed Student’s *t*-test).

## DISCUSSION

Organism survival is dependent on the intricate balance of maintaining energy homeostasis among multiple metabolically active tissues. In adults, tissue maintenance and regeneration are dependent on stem cell populations that respond to changes in physiological conditions. Nuclear receptors are transcription factors that regulate stem cell self-renewal and progeny differentiation through multiple mechanisms between tissues. Our results herein show that *ERR* is required in adult female *Drosophila* to regulate GSC lineages. Furthermore, our results show that the *Drosophila ERR* ortholog has a conserved role in regulating stem cell behavior.

Energy production in stem cells is dependent on glycolysis for maintenance of stem cell identity (Folmes and Terzic 2016). For example, embryonic stem cells are dependent on a high glycolytic flux to establish and maintain pluripotency (Tsogtbaatar *et al*. 2020), and proliferative neural progenitor cells are dependent on high levels of Hexokinase-2 to maintain glycolysis (Gershon *et al*. 2013). It was recently shown that glycolytic enzyme protein levels are higher in *Drosophila* female GSCs relative to differentiating germ cells, and loss of *Alcohol dehydrogenase* or *Enolase* results in GSC loss (Rojas-Rios *et al*. 2024). Consistent with these observations, we found that *ERR* mutant females have a reduction in Eno levels (**Figure 4**), suggesting that whole-body knockdown of *ERR* reduces glycolytic transcription in the germline. Furthermore, we found that *ERR* is required cell autonomously in the germline for GSC maintenance, however, it is likely that ERR mediates GSC self-renewal independently of glycolytic enzyme transcriptional regulation in this cell type. ERR protein is present in the cytoplasm of the germline (**Figure 1c,d**), suggesting a role for ERR outside of the nucleus. Although the *ERR::GFP.FSTF* transgene contains multiple tags that could disrupt nuclear transport, additional nuclear receptors are sequestered in the cytosol until activated (Levin and Hammes 2016). For example, Estrogen Receptor (ER) can interact with molecular chaperones, sequestering ER in the cytosol. Ligand binding triggers release of the chaperones, promoting the nuclear translocation of ER (Levin and Hammes 2016). There is currently no known ligand for ERR and it is unclear whether ERR interacts with chaperones in the germline. However, the nuclear localization of ERR in adipocytes (**Figure 5a**) but not the germline suggests that the ERR::GFP.FSTF localization in the cytosol is accurate and may suggest that there may be post-translational modifications and/or binding partners of ERR that regulate ERR nuclear translocation that remain to be identified.

In addition to glycolytic enzymes, ERR and its orthologs transcriptionally regulate members of the pentose phosphate pathway (Beebe *et al*. 2020). It was recently shown that pentose phosphate pathway enzyme transcripts are expressed in the *Drosophila* germline beginning in stage 1 egg chambers (Carvalho-Santos *et al*. 2020). This is in contrast with ERR::GFP.FTSTF, which is expressed in the entire germline. Furthermore, loss of pentose phosphate pathway activity in the germline results in death of vitellogenic egg chambers (Carvalho-Santos *et al*. 2020), in contrast to what we observed with whole-body *ERR* knockout and *ERR* mutant germline clonal analysis. Based on our results, we propose that ERR may have cytoplasmic and nuclear roles depending on the cell type; however, its cytosolic mechanism of action to maintain stem cell identity remains to be discovered.

Previous studies have highlighted adipocyte metabolism in regulating distinct processes of oogenesis in *Drosophila* (Matsuoka *et al*. 2017), including GSC self-renewal. For example, adipocyte-specific fatty acid oxidation is required for GSC maintenance, whereas, acetyl-coA production is important for early germline cyst survival (Matsuoka *et al*. 2017). Knockout of *ERR* specifically in adipocytes does not influence GSC maintenance (or any additional processes of oogenesis), suggesting that ERR activity in adult female adipocytes is not required for fertility. One possibility is that glycolysis in adipocytes is not heavily dependent on ERR activity during *Drosophila* adulthood. Consistent with this hypothesis, *ERR* activity is required during pupal stages for the induction of glycolytic gene expression, which is maintained in adults (Graveley *et al*. 2011; Beebe *et al*. 2020). In addition, enzymes involved in the TCA cycle and electron transport are globally up-regulated at the onset of adulthood in an *ERR*-independent manner (Beebe *et al*. 2020). Furthermore, downregulation of glycolytic enzymes in adult female adipocytes in response to a protein-poor diet occurs independently of changes in ERR protein levels (Matsuoka *et al*. 2017). These studies suggest that other temporal or tissue-specific regulators may act in parallel with ERR to regulate metabolism in *Drosophila* adults. In the future it will be necessary to determine the specific targets of ERR in adult versus larval adipocytes, which may provide insight to how nuclear receptor targets change under different developmental contexts.

Although ERR activity is not required in adult female adipocytes to regulate GSC number, it is possible that ERR activity is required in other somatic tissues for regulation of oogenesis. Whole-body knockout of *ERR* in adult females does decrease glycolytic and pentose phosphate pathway enzyme transcription (**Figure 2c-g**), suggesting that ERR-dependent transcription is required in at least one metabolic tissue. Single cell RNA-sequencing of adult *Drosophila* suggests moderate-to-high *ERR* transcript levels are found in the principal cells of Malpighian tubules and multiple cell types in the gut (Li *et al*. 2022), which are endocrine organs that could secrete metabolites or signaling molecules in an ERR-dependent manner to promote GSC self-renewal. In fact, it was recently shown that ERR protein is enriched in Malpighian tubule principal cells and is required for glucose-mediated glutathione regeneration and fatty acid oxidation (Holcombe and Weavers 2023). Therefore, future studies are required to dissect whether *ERR* is required in principal cells or additional cell types, as well as the downstream mechanisms utilized to regulate GSC maintenance. Furthermore, it will be of interest to determine whether the transcriptional targets of ERR are altered in a tissue-specific manner, and how these potential targets influence GSC self-renewal in adult females.

## ACKNOWLEDGEMENTS

A.B.Z., M.G.A., S.A.F., E.D.D., and L.N.W. performed experiments. L.N.W. designed experiments, analyzed and interpreted the data, and wrote the manuscript. We thank the Developmental Studies Hybridoma Bank for antibodies and the Bloomington Stock Center (National Institutes of Health P40OD018537) and the Vienna *Drosophila* Stock Center (VDRC, www.vdrc.at) for *Drosophila* stocks. *ERR* conditional knockout strains described in this manuscript were a kind gift from Katherine Beebe. We are thankful for Flybase (www.flybase.org), an essential *Drosophila* research resource. We are grateful to Heather Hundley and Jason Tennessen for critical reading of the manuscript. The data and analyses reported in this paper are described in the main figures and Supplemental Materials.

## DECLARATION OF INTERESTS

The authors declare no conflict of interests.

## FUNDING

This work was supported by the National Institutes of Health (NIH) grants R00 GM127605 (L.N.W.) and R35 GM150517 (L.N.W).

## DATA AVAILABILITY

*Drosophila* strains can be purchased from the Bloomington *Drosophila* Stock Center, the Vienna *Drosophila* Resource Center, or are available upon request if not available in the stock centers. All data generated for this study are included in the main text and figures or in the supplemental files provided.

## Notes

### Competing Interest Statement

The authors have declared no competing interest.

