## Supplementary Materials for "Estrogen-Related Receptor is Required in Adult *Drosophila* Females for Germline Stem Cell Maintenance"

**This file includes:**

Figures S1 to S7

Table S1 to S2

**SUPPLEMENTARY FIGURES**

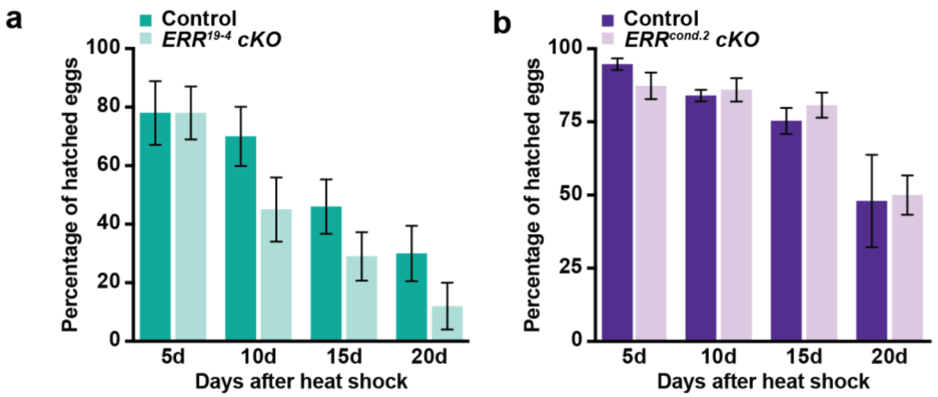


**Figure S1. *ERR* is not required in adult females for progeny survival.**

**(a,b)** The average percentage of hatched eggs of *ERR*$^{19\text{-}4}$ **(a)** and *ERR*$^{cond.2}$ **(b)** females relative to their respective controls collected from eggs laid on 5, 10, 15, and 20 days after heat shock. At least 100 eggs per genotype per timepoint were collected and observed. There were no statistical differences detected using a paired Student's *t*-test for three biological replicates.

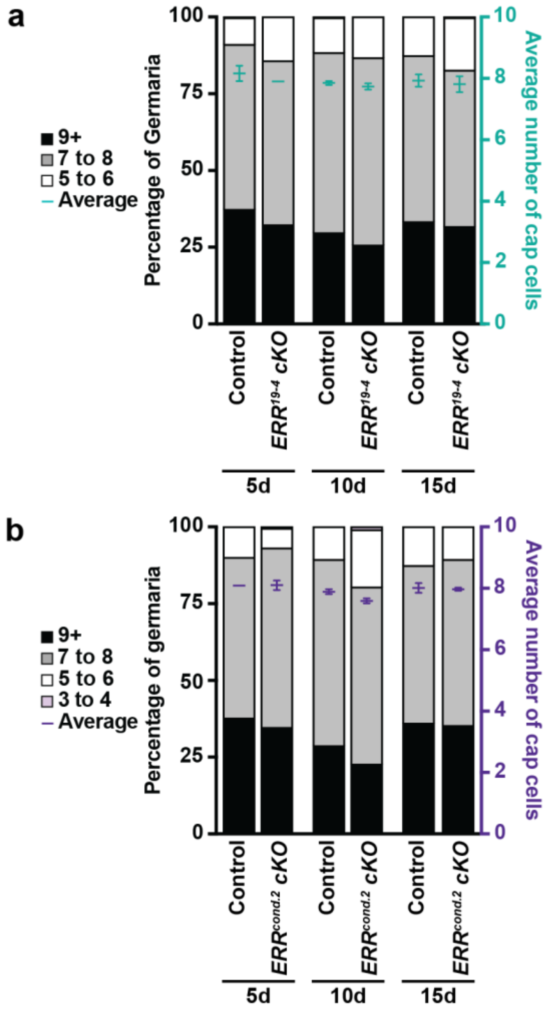


**Figure S2. Conditional knockout of *ERR* in adult females does not alter cap cell maintenance.**

**(a,b)** Bars representing the percentage of germaria containing five-to-six, seven-to-eight, or nine-or-more cap cells at 5, 10, and 15 days after the final heat shock *ERR*$^{19\text{-}4}$ **(a)** and *ERR*$^{cond.2}$ **(b)** conditional knockout females relative to their respective controls. The average cap cell numbers are shown as mean ± SEM is plotted on the right y-axis. 300 germaria were analyzed from three biological replicates. No statistical differences determined using a paired, two-tailed Student's *t*-test.

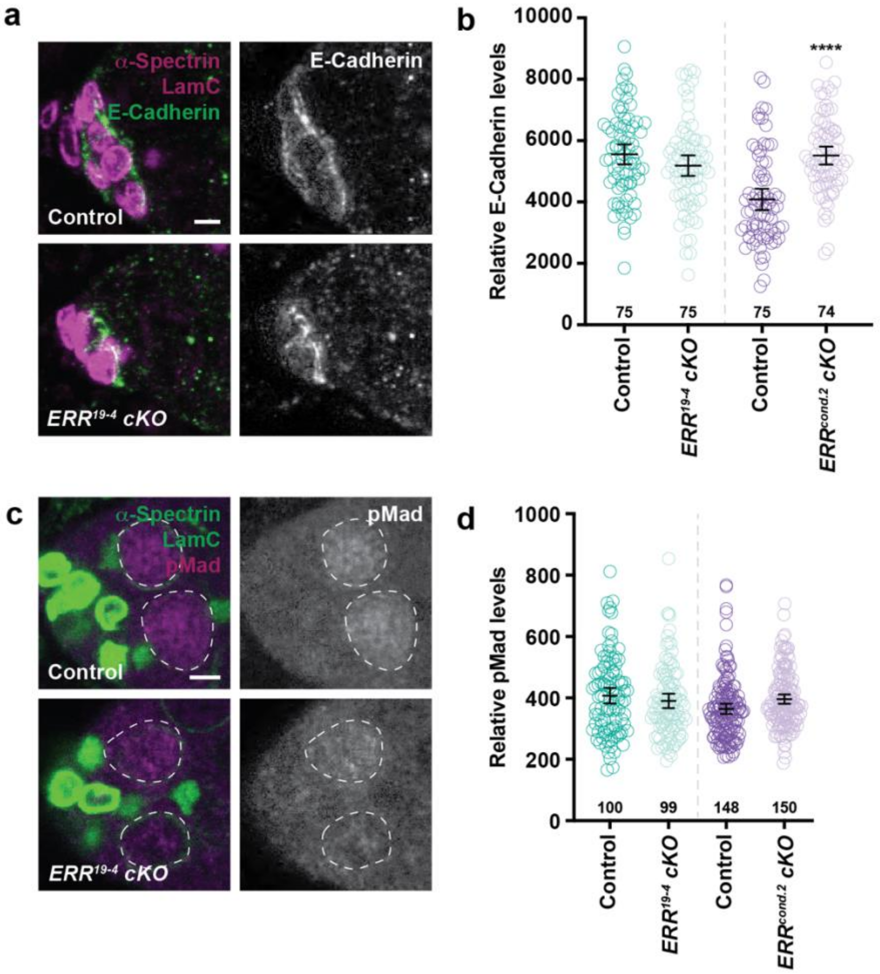


**Figure S3. *ERR* is not required in adult females for E-cadherin adhesion or BMP signaling.**

**(a)** Anterior portion of germaria from females at 10 days after heat shock from control and *ERR*$^{19\text{-}4}$ conditional knockout showing similar E-Cadherin levels at the GSC-niche junction. E-Cadherin (green); α-spectrin (magenta), fusome; LamC (magenta), nuclear lamina. **(b)** Dot plot of total cap cell E-Cadherin intensity per germarium for the experiment in (a). The number of germaria analyzed is indicated in the graph. ****$P < 0.0001$; Mann-Whitney *U*-test. **(c)** Anterior portion of germaria from control and *ERR*$^{cond.2}$ females 10 days after heat shock showing similar levels of phosphorylated Mad (pMad; a BMP signaling reporter) in GSC nuclei. pMad, BMP signaling reporter (magenta); $\alpha$-spectrin, fusome (green); LamC, nuclear lamina (green). **(d)** Dot plot of the average pMad intensity per germarium for experiment in (c). The number of analyzed GSCs for each genotype is shown in the graph. Black lines in (b) and (d) indicate mean ± 95% confidence interval for each experiment. No statistically significant differences, Mann-Whitney *U*-test. Scale bars in (a) and (c), 2.5 µm.

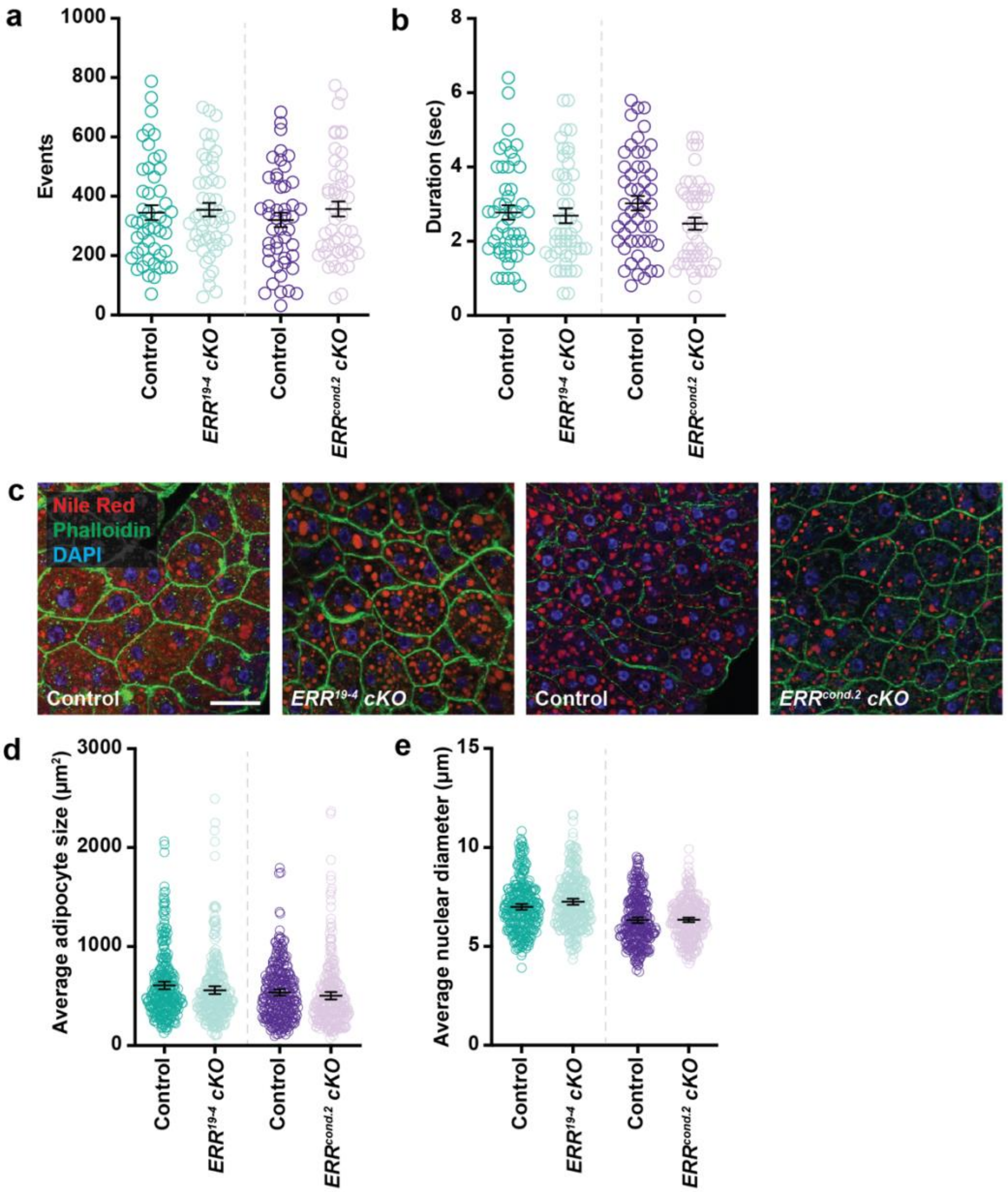


**Figure S4.** ***ERR*** **knockout in adult females does not alter feeding behavior or adipocyte morphology.**

**(a,b)** The total number of feeding events **(a)** or median time of feeding activity **(b)** for control, *ERR*$^{19\text{-}4}$, or *ERR*$^{cond.2}$ females 10 days after heat shock. Data shown as mean ± 95% confidence interval. At least 50 females per genotype were analyzed. No statistical differences, Mann-Whitney *U*-test. **(c)** Females at 10 days after heat shock in control, *ERR*$^{19\text{-}4}$, and *ERR*$^{cond.2}$

analyzed for adipocyte morphology. Adipocytes labeled with DAPI (blue), DNA; Phalloidin (green), cell membrane; and Nile Red (red), lipid droplets. Scale bar, 25 µm. **(d)** Dot plots of the average adipocyte cell area. 300 adipocytes were measured from three independent experiments. Data shown as mean ± 95% confidence interval. No statistical differences; paired, two-tailed Student's *t*-test. **(e)** Dot plots of the average diameter of adipocyte nuclei. 300 nuclei were measured from three biological replicates. Data shown as mean ± 95% confidence interval. No statistical differences; paired, two-tailed Student's *t*-test.

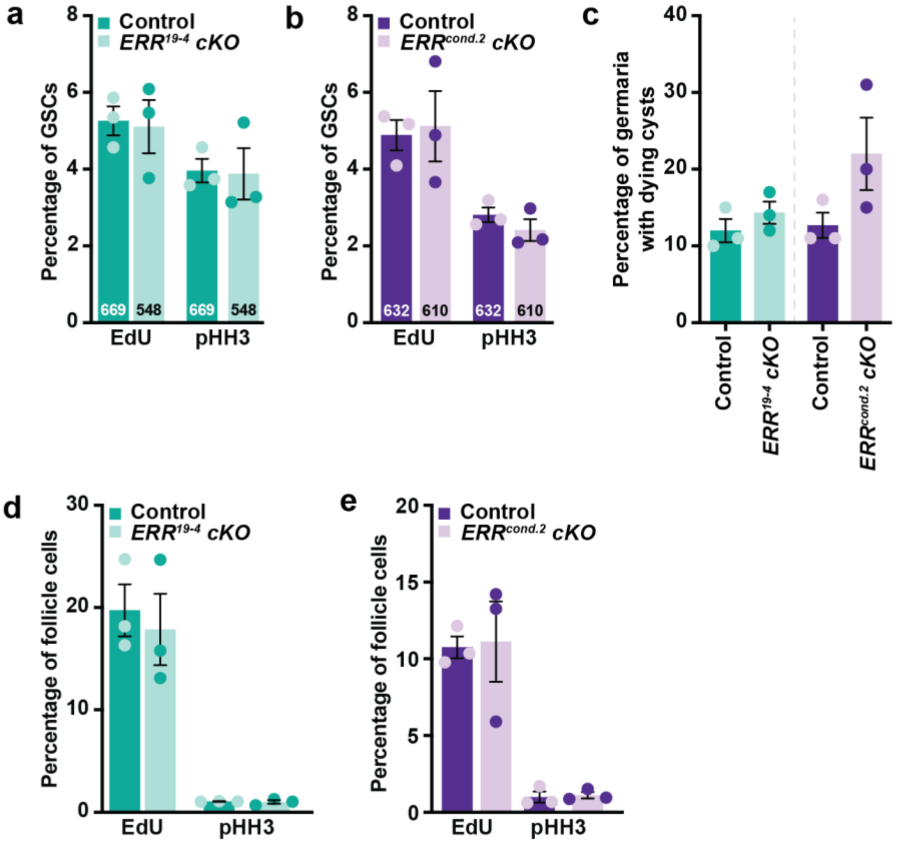


**Figure S5. GSC proliferation, early cyst death, and follicle cell proliferation do not require whole-body ERR activity in adult females.**

**(a,b)** The percentage of GSCs in S-phase (EdU incorporation) or M-phase (pHH3 labeling) in *ERR*$^{19\text{-}4}$ **(a)** or *ERR*$^{cond.2}$ **(b)** females 10 days after heat shock compared to their respective controls. The number of GSCs analyzed from three independent experiments are indicated. Data shown as mean ± SEM. No statistical differences; paired, two-tailed Student's *t*-test. **(c)** The percentage of germaria containing dying germline cysts (based on ApopTag analysis) in *ERR*$^{19\text{-}4}$ or *ERR*$^{cond.2}$ females relative to their control. 300 germaria were analyzed for each genotype. Data shown as mean ± SEM. No statistical differences; paired, two-tailed Student's *t*-test. **(d,e)** The percentage of follicle cells in S-phase or M-phase for *ERR*$^{19\text{-}4}$ **(d)** or *ERR*$^{cond.2}$ **(e)** compared to their respective controls. Follicle cells in egg chamber stages 2-6 were analyzed from 75 ovarioles (three independent experiments) and the data is shown as the mean ± SEM. No statistical differences; paired, two-tailed Student’s *t*-test.

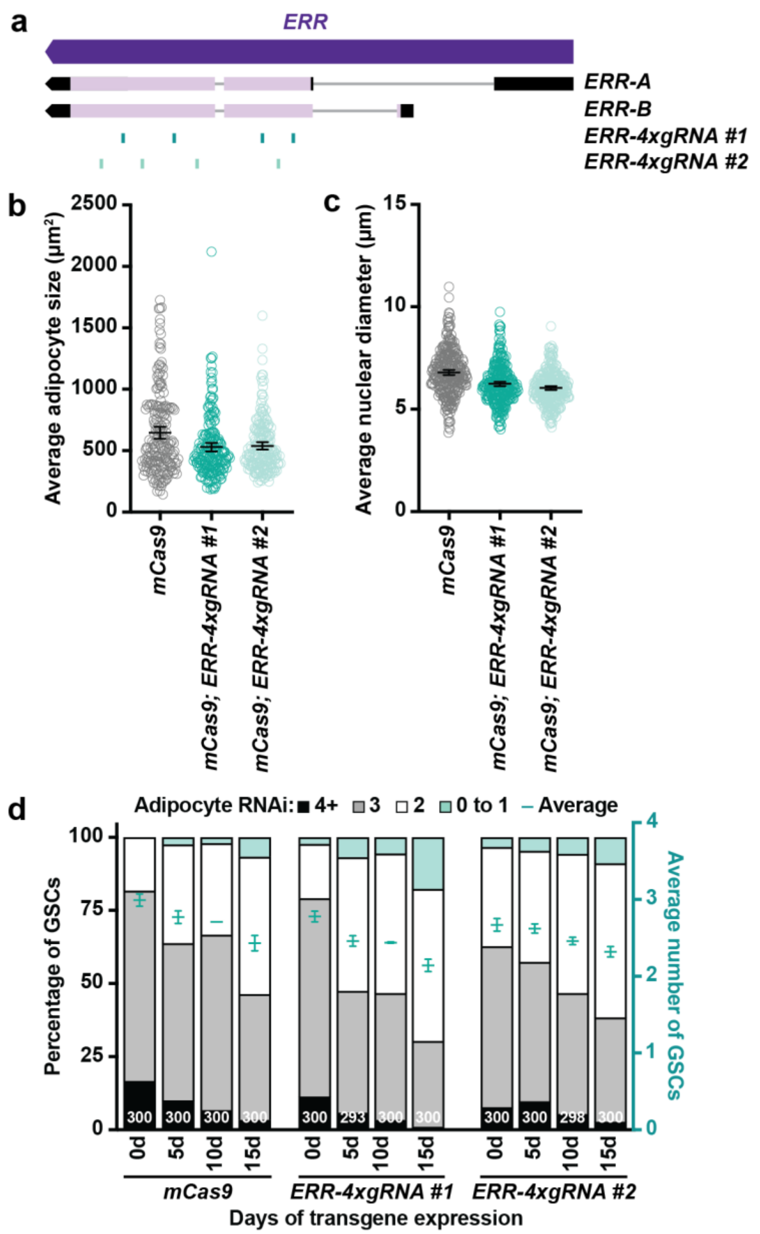


**Figure S6. *ERR* knockout in adult female adipocytes does not influence adipocyte morphology or GSC maintenance.**

**(a)** Schematic showing the *ERR* gene and its two isoforms. The sgRNA constructs (dark and light teal) *UAS-ERR-4xgRNA #1* and *UAS-ERR-4xgRNA #2* target distinct regions in the protein coding regions shared by both transcripts. **(b)** Dot plots of the average adipocyte cell area. 300 adipocytes were measured from three independent experiments. Data shown as mean ± 95% confidence interval. No statistical differences; paired, two-tailed Student's *t*-test. **(c)** Dot plots of the average diameter of adipocyte nuclei. 300 nuclei were measured from three biological replicates. Data shown as mean ± 95% confidence interval. No statistical differences; paired, two-tailed Student's *t*-test. **(d)** Bars representing the percentage of germaria containing zero-to-one, two, three, or four-or-more GSCs at different time points in control and adult adipocyte-specific *ERR* CRISPR knockout females. The average GSC numbers are shown as mean ± standard error of the mean (SEM) is plotted on the right y-axis. The number of germaria were analyzed from three biological replicates are indicated in the bars. No statistical differences, two-way ANOVA with interaction.

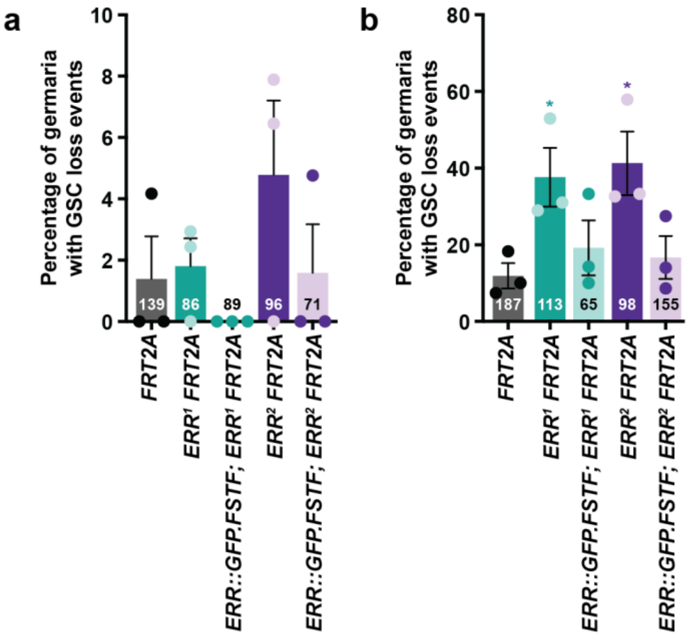


**Figure S7. *ERR* is required cell-autonomously in the germline for GSC maintenance.** The percentage of mosaic germaria with a GSC loss event at 4 days **(a)** or 12 days **(b)** after heat shock. This data is also represented in Figure 5E. The number of mosaic germaria analyzed from three biological replicates are indicated in each bar. Data is shown as mean ± SEM. $*P < 0.05$ (paired, two-tailed Student's *t*-test).

### SUPPLEMENTARY TABLES

**Table S1. Primer sequences used in this study.**

| Gene | Forward | Reverse |
|---|---|---|
| ***ERR*** | 5’-AATTGGTCAGCGTCATTGGC-3’ | 5’-TCATCTGGTCGTTAAGTGGCAG-3’ |
| ***Pfk*** | 5’-ATGGACGGATACCCATTTGC-3’ | 5’-CTTTTGCCTAATCGGAGGTG-3’ |
| ***Ldh*** | 5’-ACGGCTCCAACTTTCTGAAG-3’ | 5’-TGGGGATGATGTTCTTGAGG-3’ |
| ***Zw*** | 5’-TTTGACGGCAAGATTCCGCA-3’ | 5’-GGTAGATCTTCTTCTTGGCCAG-3’ |
| ***Pgd*** | 5’-TGCGACGAGTTAGCCAAACTT-3’ | 5’-CTGGAAGATGGGTTGGATAAGGG-3’ |
| ***Rp49*** | 5’-CAGTCGGATCGATATGCTAAGC-3’ | 5’-AATCTCCTTGCGCTTCTTGG-3’ |
| ***Act5c*** | 5’- AGGCCAACCGTGAGAAGATG-3’ | 5’- ACATACATGGCGGGTGTGTT-3’ |

**Table S2. sgRNA sequences targeting *ERR***

| ERR_4x-sgRNA #1 | ERR_4x-sgRNA #2 |
|---|---|
| CGGCCTGAAATCCTCGCCCT | AATGATGGCGATAGTCTGAA |
| GACGGATCCCGATCCAGCTC | GCTGTGCGATGTCAAGATAC |
| ATTGGGCGTAACAACGCTGG | GAAGGTCAGCTGGAGCGTCA |
| TGCGCTGTGCGATCTGGACG | CATCGTTTAGAGAATTAAGA |
